# Normative models for neuroimaging markers: Impact of model selection, sample size and evaluation criteria

**DOI:** 10.1101/2022.09.22.509002

**Authors:** Jelena Bozek, Ludovica Griffanti, Stephan Lau, Mark Jenkinson

## Abstract

Modelling population reference curves or normative modelling is increasingly used with the advent of large neuroimaging studies. In this paper we assess the performance of fitting methods from the perspective of clinical applications and investigate the influence of the sample size. Further, we evaluate linear and nonlinear models for percentile curve estimation and highlight how the bias-variance trade-off manifests in typical neuroimaging data.

We created plausible ground truth distributions of hippocampal volumes in the age range of 45 to 80 years, as an example application. Based on these distributions we repeatedly simulated samples for sizes between 50 and 50,000 data points, and for each simulated sample we fitted a range of normative models. We compared the fitted models and their variability across repetitions to the ground truth, with specific focus on the outer percentiles (1^*th*^, 5^*th*^, 10^*th*^) as these are the most clinically relevant.

Our results quantify the expected decreasing trend in variance of the volume estimates with increasing sample size. However, bias in the volume estimates only decreases a modest amount, without much improvement at large sample sizes. The uncertainty of model performance is substantial for what would often be considered large samples in a neuroimaging context and rises dramatically at the ends of the age range, where fewer data points exist. Flexible models perform better across sample sizes, especially for nonlinear ground truth.

Surprisingly large samples of several thousand data points are needed to accurately capture outlying percentiles across the age range for applications in research and clinical settings. Performance evaluation methods should assess both, bias and variance. Furthermore, extreme caution is needed when attempting to extrapolate beyond the age range included in the source dataset. To help with such evaluations of normative models we have made our code available to guide researchers developing or utilising normative models.

## 1. Introduction

Normative models, also called nomograms, are a useful tool for predicting and assessing measures for an individual relative to the population. They provide an estimate of all or part of a conditional probability distribution (often conditioned on age) for a reference population of ‘normal’ (often ‘healthy’) individuals, allowing quantification of how a certain individual might deviate from this reference population. While they are well established in some clinical contexts (e.g. paediatric growth charts (GROUP and de Onis, 2006; Borghi et al., 2006)), neuroimaging has started to adopt them only recently, where they have already been beneficial for the assessment of brain development (Erus et al., 2015; Chen et al., 2021; Dimitrova et al., 2021), ageing (Bethlehem et al., 2022; Rutherford et al., 2022) and in various clinical conditions related to psychiatry (Wolfers et al., 2018; Marquand et al., 2019; Zabihi et al., 2019; Wolfers et al., 2020) and dementia (Pinaya et al., 2021). Moving away from group-level studies, normative modelling incorporates heterogeneity in clinical cohorts, allowing predictions at an individual subject level (Marquand et al., 2016).

Compiling reference data to contextualise patient’s findings is also one of the main steps described by the quantitative neuroradiology initiative (QNI) framework for eventual adoption into routine practice of quantitative neuroradiological tools (Goodkin et al., 2019) to support clinical decision making. An example of a promising application of normative modeling of brain measures in clinical practice is in the area of dementia diagnosis (Goodkin et al., 2019). In particular, clinicians perceive information on the hippocampal volume as a valuable biomarker for cognitive impairment evaluation in suspected Alzheimer’s disease patients (Bosco et al., 2017). Thus, computation of normative curves for hippocampal volume for an early assessment of disease onset could provide valuable assistance to clinicians.

One of the drivers for the recent increase in the development and use of normative models is the availability of big data. Studies that collect huge amounts of data, such as ADNI (Jack et al., 2008), ABIDE (Martino et al., 2014), dHCP (Hughes et al., 2017; Makropoulos et al., 2018), HCP (Van Essen et al., 20134), UK Biobank (UKB) (Miller et al., 2016), make it possible to conduct big population studies and apply normative models to brain MR imaging data. Generally, having a large sample size allows more precise normative distributions to be calculated. However, it is not known what is the minimal sample size for providing a realistic and sufficiently accurate normative model for neuroimaging applications. This is important as even though neuroimaging datasets are now much larger than before, they are still a lot smaller than other medical datasets that have been used to create normative models in other areas (e.g. paediatric growth charts).

The development and choice of the best model to generate a normative model for a particular application is crucial and a very active area of research. Currently available normative modelling methods use a range of techniques, including hierarchical linear models, polynomial regression, quantile regression, support vector regression and Gaussian process regression (Marquand et al., 2019). However, each technique has some weaknesses, for example, Gaussian process regression does not scale well with dataset size, linear models do not capture nonlinear relationships, and other methods make assumptions of Gaussianity about the conditional distribution (i.e. a consistent and symmetric relationship between all the percentile curves). One good candidate for more flexible modelling is the generalized additive model, often implemented using the VGAM (Vector Generalized Linear and Additive Models) (Yee, 2015) or GAMLSS (Generalized Additive Models for Location Scale and Shape) (Rigby and Stasinopoulos, 2005) packages.

GAMLSS is a very flexible unifying framework for univariate regression (Stasinopoulos et al., 2017) that accommodates a wide range of distribution models where all the parameters of the distribution can be modelled as a function of the explanatory variables. It therefore extends basic statistical models, allowing flexible modelling of non-constant variance, skewness and kurtosis in the data.

There are many examples of GAMLSS being used in practice for normative modelling in neuroimaging, with a range of training sample sizes; for example, 94 fetal images in Ber et al. (2017), 948 pediatric images in Dong et al. (2020), 19,793 adult images in Nobis et al. (2019), 25,575 pediatric and adult images in Córdova-Palomera et al. (2021), and up to 123,984 images across the majority of the lifespan in Bethlehem et al. (2022). This demonstrates the wide range of training sizes used, but despite this the issue of whether the number of images used for training is sufficiently accurate, and quantitative measures of accuracy, are rarely discussed.

Alternative methods to GAMLSS have also been used in neuroimaging and include the lambda-mu-sigma (LMS) method and implementations in VGAM (Schmidt-Richberg et al., 2016; Vinke et al., 2019; Vernooij et al., 2018) (with training sizes of 248 through to 4915 in these papers), warped Bayesian linear regression (Fraza et al., 2021) (with 20,083 adult images), as well as sliding window approaches with both fixed and variable window sizes, such as Nobis et al. (2019) (with 19,793 adult images), and Janahi et al. (2021) (with 40,000 adult images).

A common factor across many implementations is the use of a transformation function, such as affine, Box-Cox and Sinh-Arcsinh (SHASH), applied to a standard normal distribution. Two notable works that have investigated the merits of different transformations (Fraza et al., 2021; Dinga et al., 2021) both concluded that SHASH was the best transformation, with the latter paper using GAMLSS. We will therefore show many results from GAMLSS with SHASH in our results, although we tested several alternatives as well.

Evaluation of the normative model obtained in most studies in neuroimaging to date is either missing or not performed in ways that address the key needs of the intended application. For example, in clinical settings it is common to estimate the outer percentile curves (e.g., the 5^*th*^ percentile), to identify participants that are most likely to have some disease. Assessing errors in the central tendency or explained variance (Rutherford et al., 2021) does not provide crucial information about the accuracy of key percentile estimates. Furthermore, providing breakdowns of errors so that the errors at the ends of the distribution can be assessed is important given that, in real life applications, the density of data points typically decreases at one or both ends of the age range in the sample. Knowing the behaviour at the edges is important for setting the practical limits of the normative model and estimating the likely performance of any extrapolation. Another factor that needs to be considered in evaluation is whether bias can be assessed, which is difficult to accurately do without ground truth or massive amounts of data.

In this study we evaluate the effect of sample size and model selection on normative models for neuroimaging markers, using hippocampal volume as an exemplar to help with the narrative but without this in any way limiting the scope of our investigations. Our approach is to use a range of simulated data from a known ground truth and can therefore assess bias and variance in any of the percentile curves. From this we aim to provide guidelines on choosing appropriate sample sizes and fitting methods with respect to the levels of bias and variance that can be expected. The investigations focus on the outer percentiles (1^*st*^, 5^*th*^ and 10^*th*^ percentiles), as these are the most clinically relevant.

A range of models are included, but our goal is not to perform an exhaustive search to find the best possible model, and so a number of commonly applied modelling methods are included (e.g. GAMLSS and sliding windows), which are intended to be representative of the range of methods used in practice. Our approach can easily be used to evaluate the performance of any normative model. It could therefore be used as a power calculation tool to assess the expected variance and bias of the percentiles of interest for particular normative modelling studies and applications.

## 2. Methods

An overview of the workflow for one of the normative modelling methods is presented in Figure 1. We initially select an analytical ground truth distribution for a scalar quantity (what we will refer to as hippocampal volumes as an exemplar) in the age range of 45 to 80 years (to approximately match the UK Biobank study). This distribution is then used to create many simulated samples and the normative modelling method is to fit to each simulated sample. Multiple sets of simulated data were generated for each sample size (where sample sizes vary from 50 to 50,000) so that for each model, at each sample size, there were many fits. Finally, we evaluated how close the fitted percentile curves are to the ground truth curves, concentrating on the 1^*st*^, 5^*th*^ and 10^*th*^ percentiles as these have the most clinical utility, given that the 90^*th*^, 95^*th*^ and 99^*th*^ percentiles should follow the same patterns.

**Figure 1:**
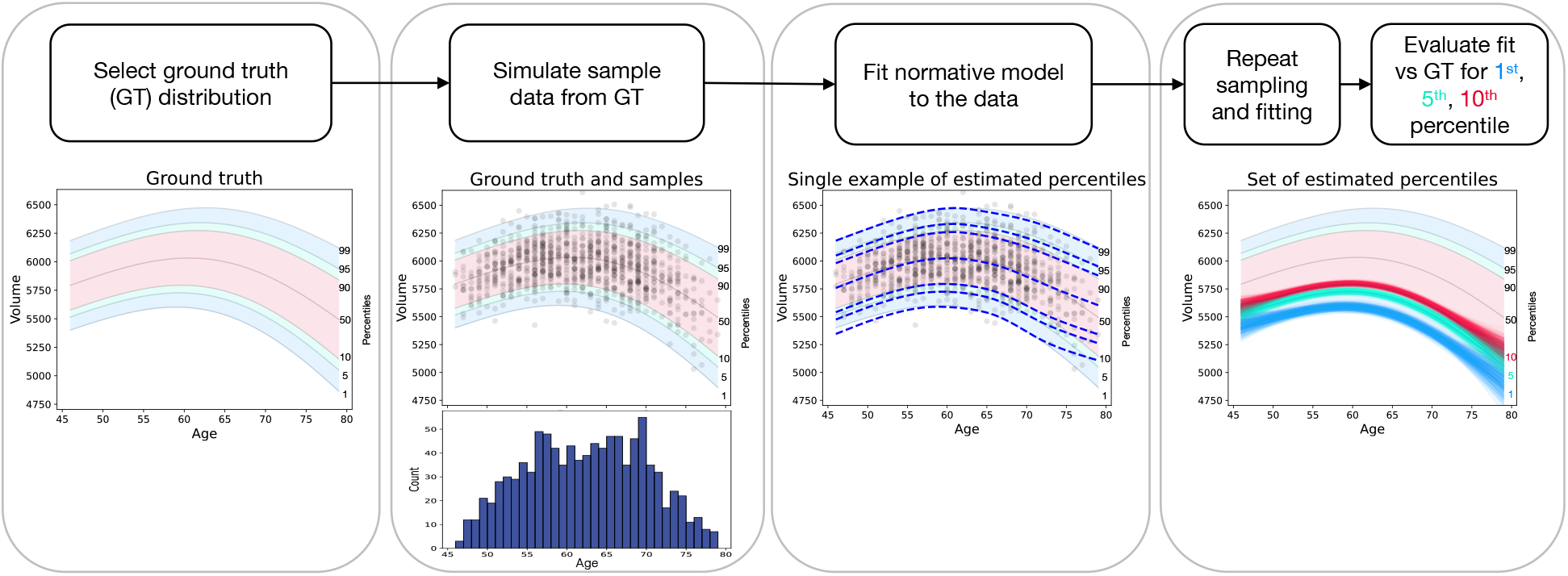
Overview of the framework for simulating and fitting a normative modelling method. The diagram summarises the methodological steps performed in the present study. For each step, an example result is given for a simulated sample size of *N*_*s*_ = 1, 000 and one modelling method. For more details on the different options tested for each step, please refer to the main text.

### 2.1. Simulated samples

We generated samples with the following sizes: 50, 100, 200, 500, 1,000, 2,000, 5,000 and 50,000, each data point/participant corresponding to one scalar value (hippocampal volume) within an age range 45 to 80 years. For each sample size, *N*_*s*_, we randomly generated *N*_*D*_ simulated samples (i.e. a set of hippocampal volumes) where *N*_*D*_ was 1,000 except for *N*_*s*_ = 50, 000. In that case we set *N*_*D*_ to 100 since, unsurprisingly, the variation in the results was a lot less and we considered that the additional data transfer and computational times were not justified.

Two ground truth distributions were chosen to cover the simplest possible case and a slightly more difficult, but plausible, case: (i) linear mean and constant variance (LinMean - ConstVar), and (ii) nonlinear mean and non-constant variance (NonLinMean NonConstVar). More precisely, the two ground truth functions were:

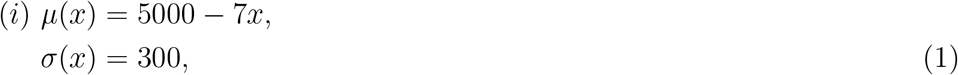

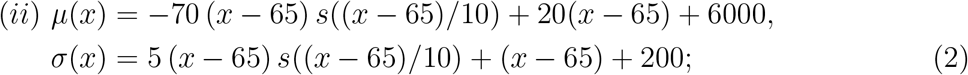

where *x* is age in years and *s*(*x*) = 1*/*(1+exp(−*x*)) is the sigmoid function. In the latter case *μ*(65) = 6000, *σ*(65) = 200 and because the sigmoid is near zero for large negative values and near one for large positive values, it smoothly interpolates between the asymptotic lines *μ*(*x*) = 20(*x* − 65) + 6000 and *μ*(*x*) = −50(*x* −65) + 6000, where the scaling of (*x* − 65) inside the sigmoid controls how rapid the transition is between these.

In both cases the conditional distribution, *p*(*y*❘*x*) (i.e. for hippocampal volume at a fixed age) is Gaussian, with mean of *μ* and standard deviation of *σ*. Non-Gaussian distributions could easily be incorporated into the simulations, but we were more interested in how well the normative models would perform in relatively simple circumstances, given that we do not know how non-Gaussian something like hippocampal volume is likely to be in reality. Making the true distributions slightly simplified also gives us an estimate of the upper limit of the performance of the normative modelling methods.

The distribution of ages was not uniform but chosen to be approximately similar to the UK Biobank project (Miller et al., 2016) distribution, as this shows the typical characteristic of having fewer participants at the ends of the age range (see one example at the bottom of the second panel in Figure 1 and supplementary Figure S1 for other sample sizes). For each sample size a set of ages was chosen randomly from this distribution but then fixed for all simulated samples with that size, so that the values of interest (e.g. hippocampal volumes) were varied but the participant ages were not, to limit the variation being tested to one variable only. This is again a slightly simplified setting, as if both were to vary then the results obtained would likely be even more variable.

### 2.2. Normative models and fitting methods

We used fitting methods based on sliding windows and generalised additive models for location, scale and shape (GAMLSS).

A sliding window method is a model-free analysis in which an age-window of a variable or fixed size is moved along the age axis, calculating a summary quantity (e.g. average or percentile) of all values falling within the window. We have implemented two versions: (i) a fixed age window of size 5 years (SliWinW5); and (ii) a variable window where the size is adjusted to include 10% of the participants for each centre position (SliWinP10), which matches Nobis et al. (2019). The latter case has a potential advantage at the lower and higher ends of the age range where the number of participants is more sparse, as this would then adjust the size to include a wider age range, although it also might over-regularise the curves by doing this. In both cases the result was then slightly smoothed using a Gaussian kernel with full width half maximum (FWHM) of 5 years.

GAMLSS is a four parameter distribution, modelling *μ, σ, ν* and *τ*, which are shape parameters of the distribution related to the mean, variance, skewness and kurtosis of the distribution. The implementation we used for the gamlss function came from package GAMLSS (version 5.3-4) (Stasinopoulos et al., 2017). We used several GAMLSS models, implementing linear fitting or cubic spline smoothing across age, together with a Box Cox T (BCT) (Rigby and Stasinopoulos, 2006) or a SinhArcsinh (SHASH) (Jones, 2005) transformation, both of which create four parameter continuous distributions.

More specifically, we used the following models: (i) linear fitting (denoted as BCT-linear or SHASH-linear); (ii) cubic spline smoothing where only one parameter, *μ*, was modelled as a function of age (denoted as BCT-*μ* and SHASH-*μ*); and (iii) cubic spline smoothing where two parameters were modelled as a function of age, namely location *μ* and scale *σ*, (denoted as BCT-*μ*-*σ* and SHASH-*μ*-*σ*). The remaining parameters (including skewness and kurtosis) were kept constant with respect to age but were estimated from the data and not dependent on other variables, following the conclusions of Dinga et al. (2021). They showed minimal differences between the predictions from simpler models with only age-dependence for location and scale (*μ* and *σ*) and the predictions from more complex models with all four parameters depending on age.

### 2.3. Evaluation

As the simulations are based on known ground truth, both bias and variance can be assessed. This was one of the main reasons for conducting this study. We were also primarily interested in the ability to model the outer, clinically-relevant percentiles. Consequently, we used two main types of evaluation: (i) comparisons of the model percentile curves with the ground truth curves in terms of difference in the principal quantity of interest (e.g. hippocampal volume), and (ii) calculation of the percentage of the ground truth distribution falling beneath a model’s estimated percentile curve (i.e. what the true percentile is for each point on the estimated curve). In each case we assessed both the bias and variance by using signed errors, in volume or percentile values. Central values (median, mean, etc.) measure the bias (i.e. consistent offsets from the true value) and the width of the distribution (IQR, standard deviation, etc.) measures the variance or variability in the estimates.

In common practical settings the variance can be measured easily (e.g., measuring variation in cross validation methods across different folds) but it is much more difficult to assess bias without knowing the ground truth. The percentage of test set data points beneath a model curve can be estimated from real data when the ground truth is not known, but this requires extremely large samples for accurate estimation.

For example, using the binomial distribution for binned data points would give a standard deviation for a p-value estimate (i.e., a percentile) of 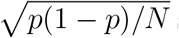 such that for *p* = 0.01 this requires *N*_*s*_ = 396 (within a single bin) to obtain an estimate of 0.01 *±* 0.005, or *N*_*s*_ = 2475 to reduce it to 0.01 *±* 0.002. More sophisticated evaluation estimation methods can improve a little on this, but it clearly shows the order of magnitude required, which demonstrates that extremely large test set sizes are required for accurate evaluations.

The ground truth distribution is defined by the conditional probability density *g*(*y* ❘*x*), where *y* is the hippocampal volume and *x* is the age. From this the associated cumulative distribution (along *y*) can be defined as 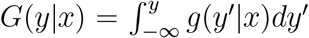. A percentile curve, for a fixed percentile value *p*, is then implicitly defined as the points where *G*(*y*❘*x*) = *p*; i.e., a curve with respect to *x*, for fixed *p*, given by *y*_*g*_(*x, p*) where *G*(*y*_*g*_(*x, p*) ❘*x*) = *p*.

When a particular sample of size *N*_*s*_ has been fit by a normative model, it either explicitly or implicitly defines an estimated conditional density *f* (*y*|*x*), along with the associated percentile curves *y*_*f*_ (*x, p*).

The two performance measures that we will use are:

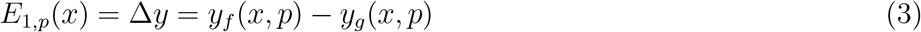

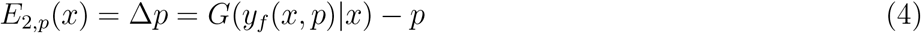

where *E*_1,*p*_ is the difference between the estimated and ground truth percentile curves, in units of hippocampal volume, and *E*_2,*p*_ is the difference between the true percentile, *p* and the percentile at the estimated value *y*_*f*_ (*x*; *p*), where *G*(*σ*) maps the volume value to the cumulative probability (i.e. the percentage of the ground truth density under this value).

When summarising the distribution of these error values over instances of simulated data we use: (i) 95% range of volumes for *E*_1_(*x*) (calculated as the 97.5^*th*^ percentile - 2.5^*th*^ percentile of the *E*_1_(*x*) values across simulations); and (ii) mean absolute error (MAE) for *E*_2_(*x*) (i.e. mean(❘*E*_2_(*x*) ❘)). When values are summarised over ages, the value at each age is counted equally (regardless of how many data points exist with that age) for calculating the mean. Note that we never summarise results across different percentiles, and whenever the *p* index is missing on the error (e.g. *E*_1_ and *E*_2_) it should be considered to be there implicitly.

### 2.4 Summary

The different options being explored here are outlined in Table 1. This shows that there are 5 different options, with anywhere from 2 to 8 possible settings, leading to a large number of different combinations to explore. Results for all combinations were generated, but these will be presented in a systematic way, keeping certain options fixed or summarising over the different settings, so that the most important aspects are clearly laid out.

**Table 1.**
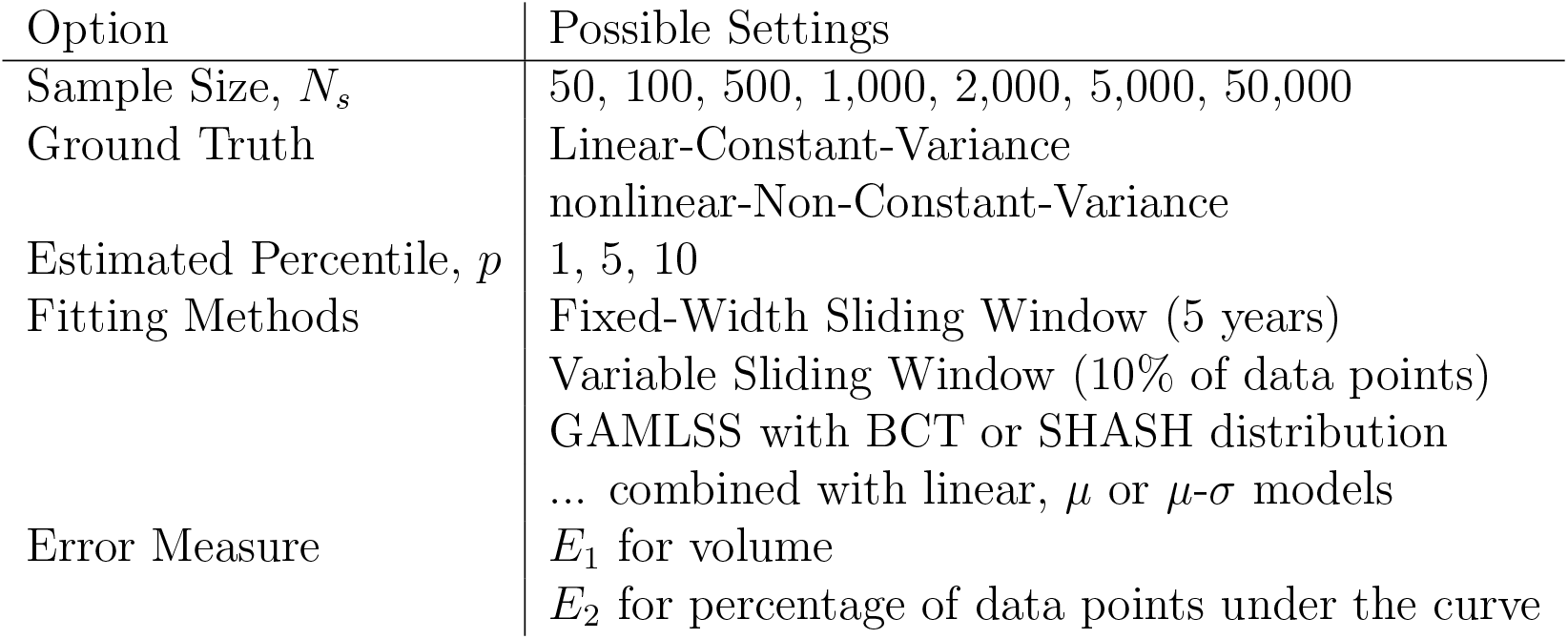
Summary of the different options explored in this work.

## 3. Results

The following sections will present results dissected in different ways, starting by looking at the effects of sample size and fitting method in general. Following this is a more detailed look at how the results vary with age, across different percentiles and sample sizes. The last result then more explicitly investigates the bias and variance components, with particular emphasis on their relation with age and sample size.

### 3.1. Sample size

Figure 2 shows the effect of sample size on the errors in the volume estimates, *E*_1_. The 95% interval is used to purely capture the variance of the error, independent of bias, and is shown for the first and fifth percentiles (columns) with different ground truth functions (rows). The values are calculated for each age, across all the simulated samples at a given sample size, and then the mean across the age range is taken, equally weighting each age. Results for 10^*th*^ percentile are similar and can be found in the supplementary material.

**Figure 2:**
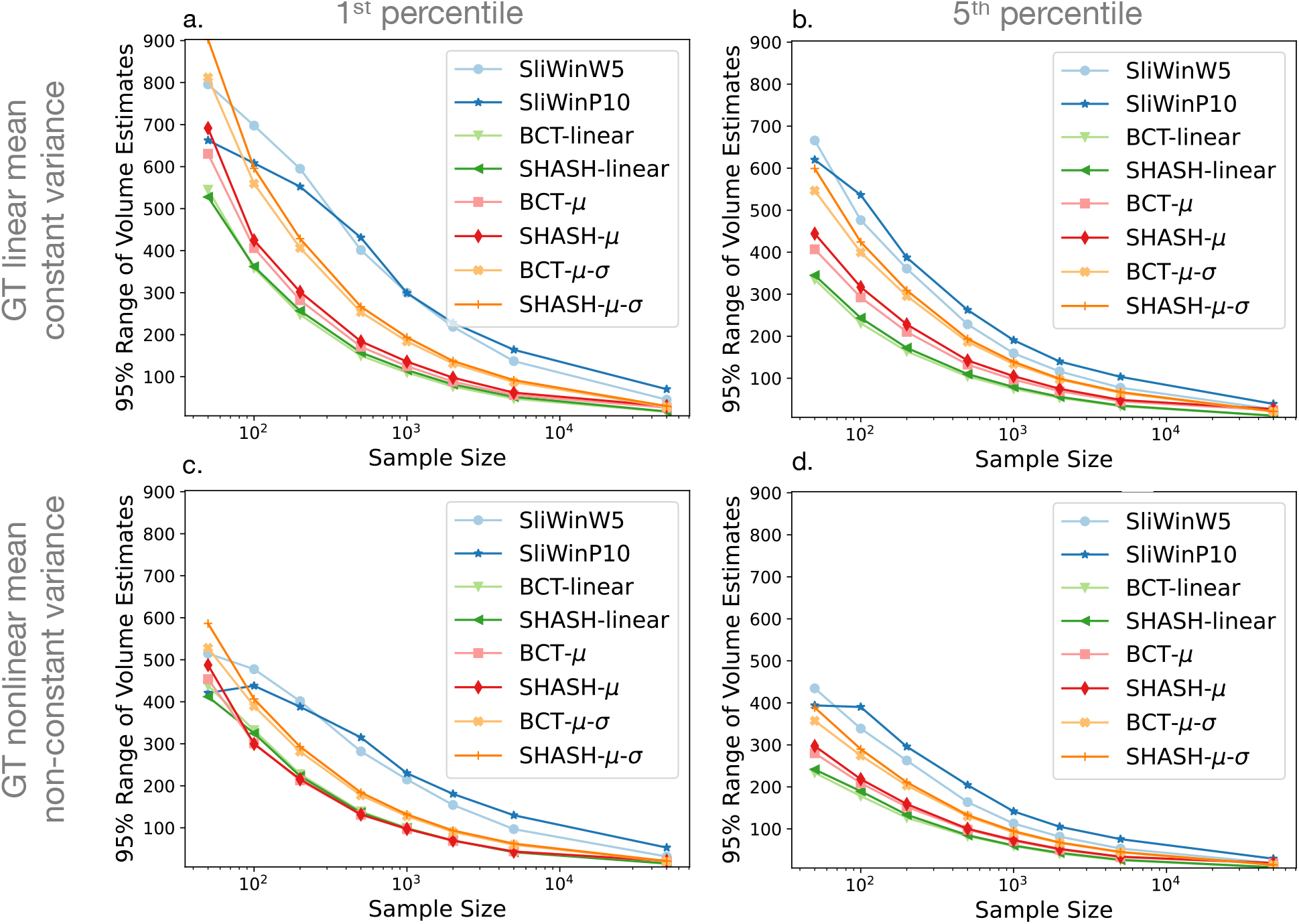
95% range of volume estimates, *E*_1_, against sample size for each fitting method. The plots show the mean across age of the 95% intervals of volume errors (*E*_1_) of the 1^*st*^ percentile (left column) and 5^*th*^ percentile (right column) curves for linear mean and constant variance ground truth (top row) and nonlinear mean and non-constant variance ground truth (bottom row). Results for the 10^*th*^ percentile were very similar to those for the 5^*th*^ percentile and are shown in supplementary figure S2.

Similarly, Figure 3 shows the effect of sample size on the errors in percentile value, *E*_2_.

**Figure 3:**
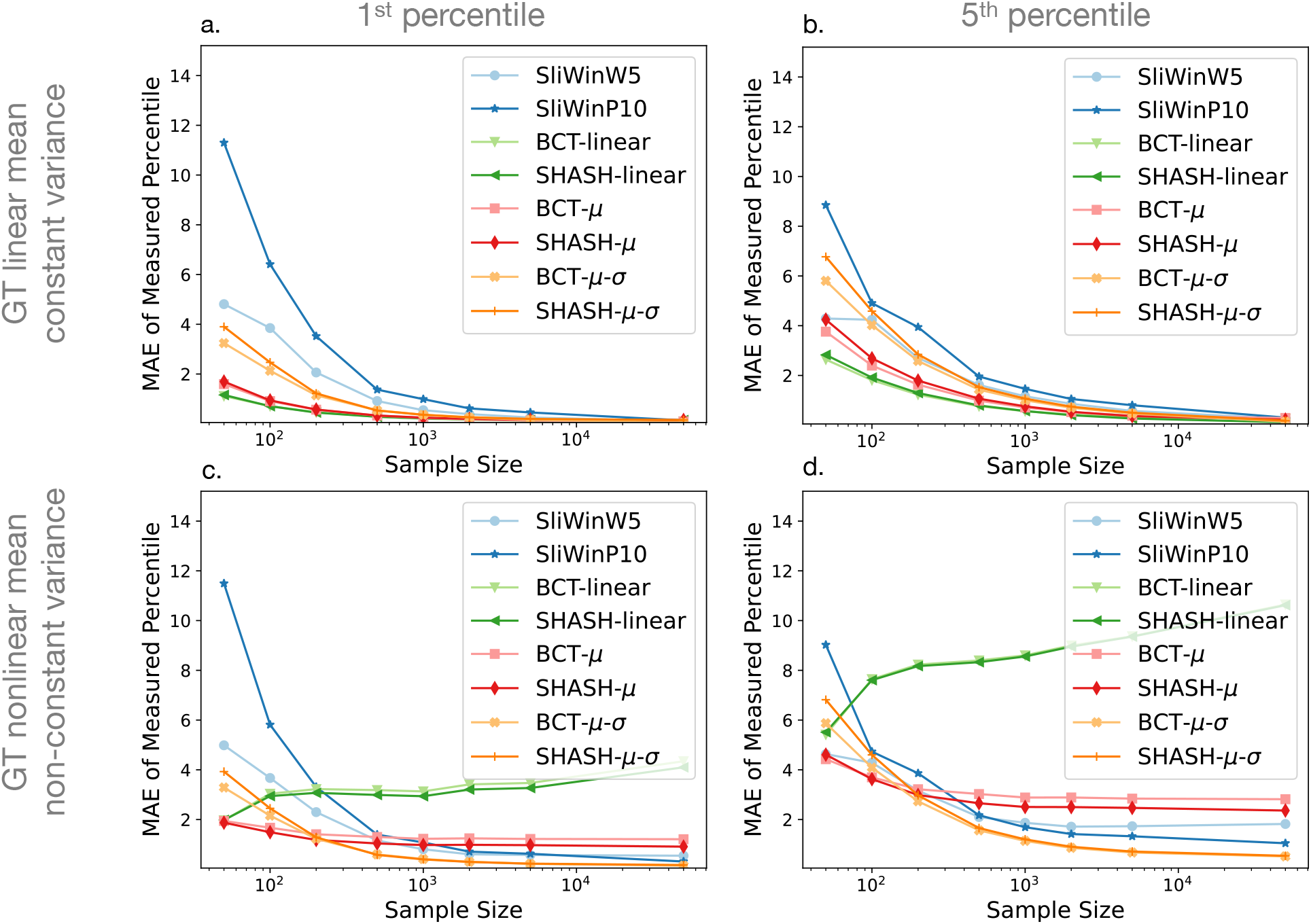
Mean Absolute Error (MAE) of *E*_2_ against sample size for each fitting method. The plots show the mean across age of the mean absolute error of the estimates of percentile error (*E*_2_) of the 1^*st*^ percentile (left column) and 5^*th*^ percentile (right column) curves for linear mean and constant variance ground truth (top row) and nonlinear mean and non-constant variance ground truth (bottom row). Results for the 10^*th*^ percentile were very similar to those for the 5^*th*^ percentile and are shown in supplementary figure S3.

The mean absolute error (MAE) in this case reflects both bias and variance in the error, and is used partly because of the floor effect in *E*_2_, as a percentile cannot be less than zero. The same type of limit does not apply in the other direction. e.g. errors can easily be 5% or more on one side. Note that in the tail of the distribution the difference in behaviour between the two error estimates, *E*_1_ and *E*_2_, becomes greater, because a small change in the percentile can correspond to a large change in volume, whereas for higher percentiles a large change in the percentile typically corresponds to only a small change in volume. As in Figure 2, the results in Figure 3 present results for the first and fifth percentiles as columns with different ground truth functions presented in rows, with results for the 10^*th*^ percentile curves reported in the supplementary material.

For all fitting methods and ground truth functions, the errors decrease with increasing sample size, as expected. The MAE of the percentiles, *E*_2_, also initially decreases with larger sample size, but there is not much improvement at very large N, especially in the case of the nonlinear ground truth (Figure 3 panels c,d). Nonlinear methods show comparable performance to linear methods in the case where the ground truth function is actually linear, but unsurprisingly the linear methods perform poorly in the case of a nonlinear ground truth. The differences between individual fitting methods are relatively minor for the most part.

However, the sliding window methods nearly always perform weaker than the GAMLSS methods. A closer look at Figures 2 and 3 indicates that, amongst the GAMLSS variants, the one with the SHASH-*μ*-*σ* transformation performs better, for both error measurements, which is most evident when assessing the fifth percentile with nonlinear ground truth (panels d in both figures). An apparent discrepancy in the sliding window results in this case can be seen, where they are clearly the worst in Figure 2d but third/fourth best in Figure 3d, for high N, which can be explained by the fact that large underestimates in volume are typically associated with small changes in percentile errors. Since these results accord with results from the literature showing superior performance for the GAMLSS method with the SHASH-*μ*-*σ* transformation, we will only show results for this method in further analyses.

### 3.2. Uncertainty across age

The results presented so far show the performance for each fitting method with different sample sizes and methods, but summarising over all ages. Figure 4 illustrates performance as a function of age for both a single example (panels a1, b1, c1 and d1) and summarising over all samples of this size (panels a2, b2, c2 and d2), with a fixed sample size of *N*_*s*_ = 1, 000 in all cases. The four different pairs of panels show results when varying the ground truth (top and bottom rows for linear and nonlinear respectively) or the fitting method (left and right columns for GAMLSS SHASH-linear and GAMLSS SHASH-*μ*-*σ* respectively).

**Figure 4:**
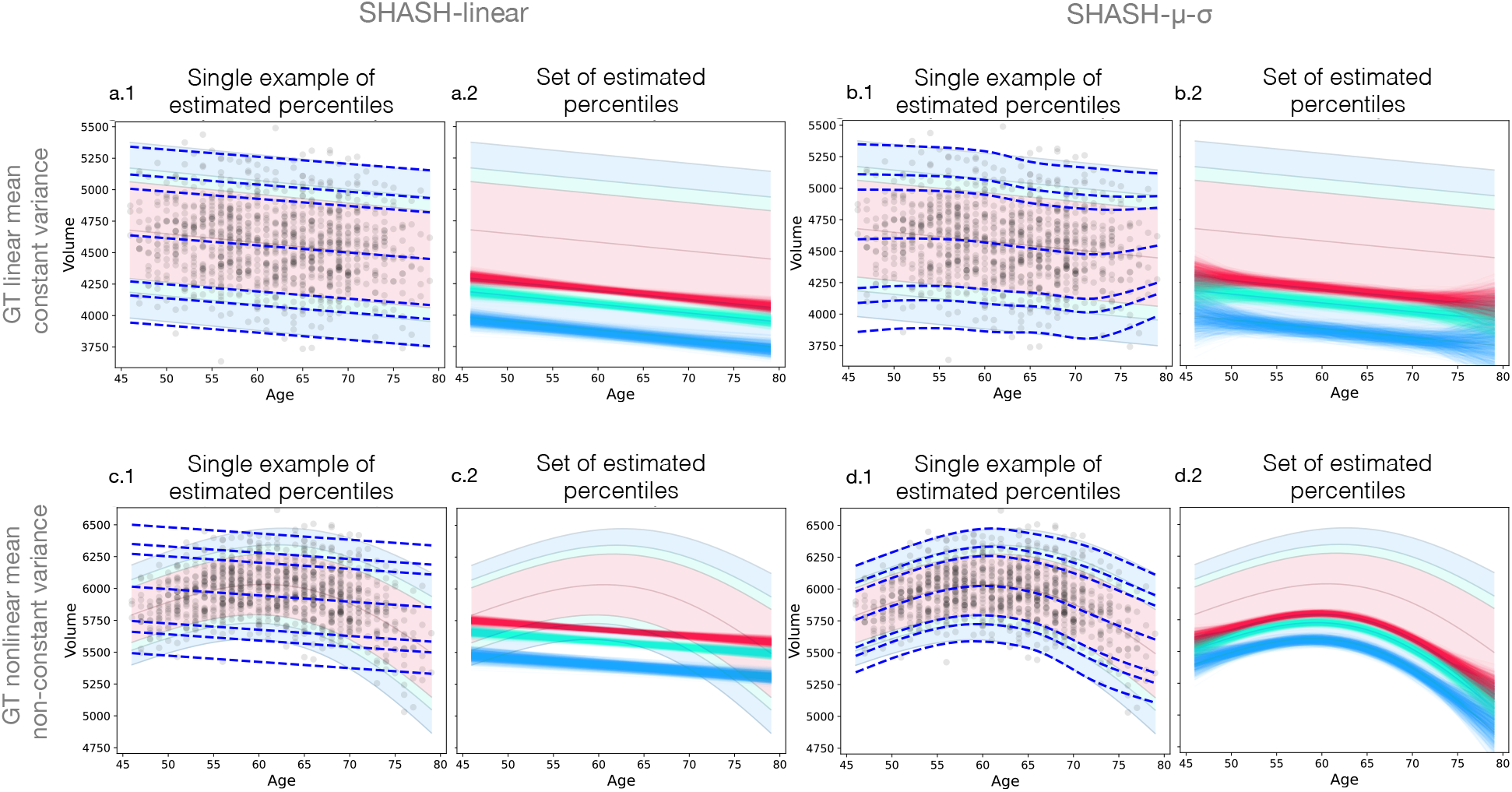
Examples of estimated percentile curves for SHASH-linear (left) and SHASH-*μ*-*σ* (right) and ground truth simulation functions (rows) using sample size 1,000. Panels are grouped in pairs with the left ones (a1, b1, c1, d1) showing a single example of a simulated sample (black circles) with estimated percentiles (blue dashed lines) overlaid onto the ground truth percentiles of 1, 5, 10, 50, 90, 95 and 99. Panels on the right in each pair (a2, b2, c2, d2) show the 1^*st*^ (blue), 5^*th*^ (green) and 10^*th*^ (red) percentile curves of all simulated samples.

As expected, the results are extremely poor for the linear fitting method when applied to the nonlinear ground truth (c2). But the linear fitting method is superior, especially at the ends of the age range, when the ground truth is linear (b2 vs a2). The nonlinear fitting method is slightly worse than the linear version when the ground truth is linear, but it nonetheless provides reasonable fits except for the ends of the age range. It provides a vastly better fit for the nonlinear ground truth (d2), although again it is less accurate near the ends of the age range. Note that even when the fit is very poor, such as in panel c2, the variance of the estimated percentile curves may be small, even though the bias is very large.

It is important to note that these simulated samples have a smaller number of data points at the ends of the age range. This reflects the properties of many real datasets, e.g. the UK Biobank imaging study, which was used as a basis for our age distribution (see example histogram in Figure 1). The rapid increase in errors *E*_1_ or *E*_2_ at the end of the age range (Figures 2 and 3) highlights the fact that summary measures over age may hide information that could be extremely important in a number of applications.

Figure 5 shows how sample size affects the estimated percentile curves across the age range using GAMLSS with SHASH-*μ*-*σ* with linear and nonlinear ground truths. It can be seen that the variability substantially decreases as the sample size gets larger, although higher variability, as well as bias, can still be observed at the ends of the age range. For example, the 5^*th*^ percentile curve at the high end of the age range for the nonlinear ground truth shows not only variance but noticeable bias. The sample size varies over three orders of magnitude here, and it is only for the largest sample of 50,000 data points that the variability becomes narrow compared to the spacing of the percentile curves.

**Figure 5:**
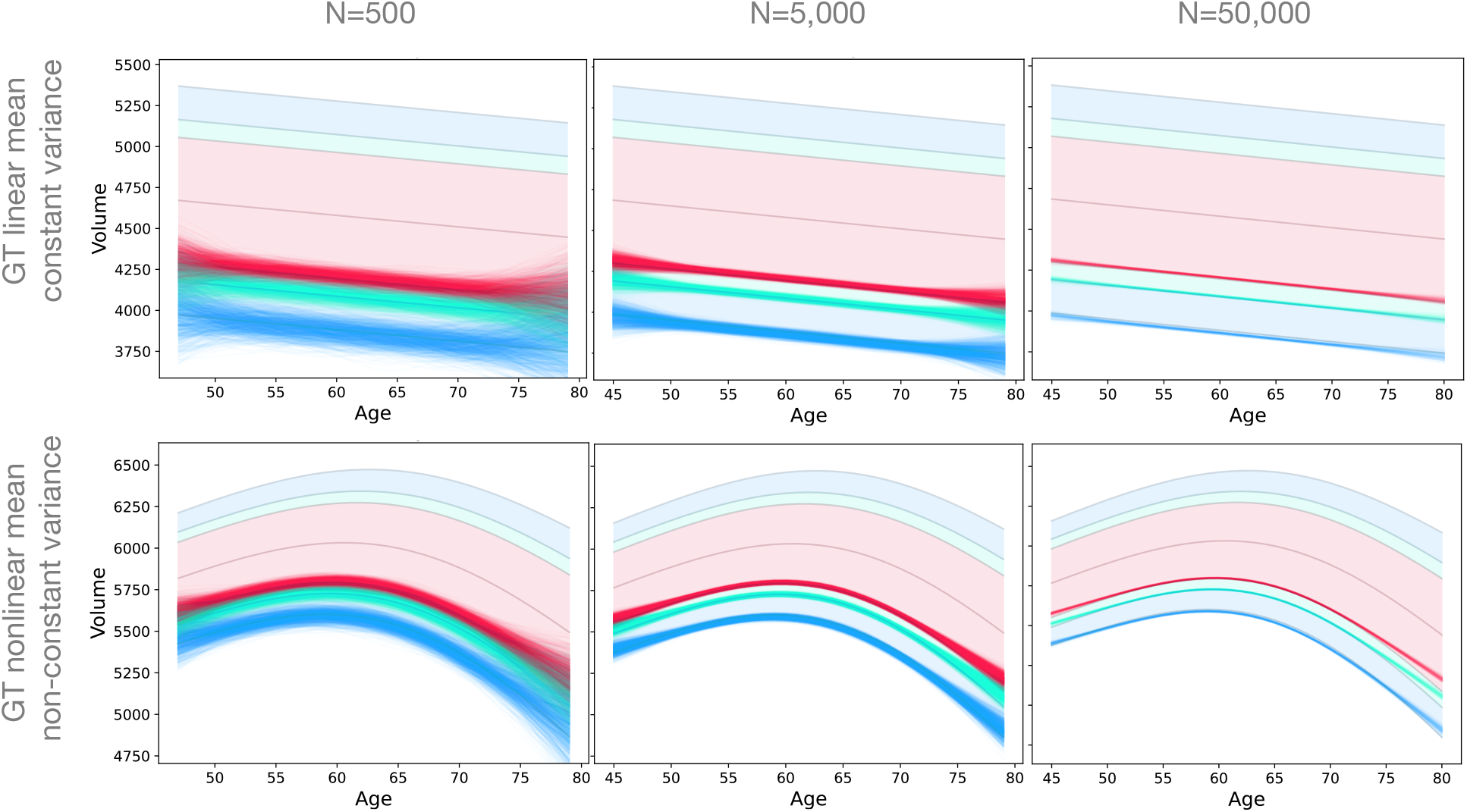
Examples of estimated 1^*st*^ (blue), 5^*th*^ (green) and 10^*th*^ (red) percentile for sample sizes (columns) of 500, 5,000 and 50,000 and ground truth simulation functions (rows) using the SHASH-*μ*-*σ* model.

A more quantitative analysis for performance with respect to age is shown in Figure 6 for the nonlinear ground truth and GAMLSS SHASH-*μ*-*σ* fitting method. Here the results are separated into bias quantified by the median error for *E*_1_ (top row), and variance components quantified by the interquartile range of *E*_1_ (middle row) for the 1^*st*^ (blue), 5^*th*^ (green) and 10^*th*^ (red) percentiles. These values are in units of volume and it is helpful to keep in mind that the interquartile range in the ground truth is approximately 500 in these units. It can be seen from the figure that the bias does not change very much across sample sizes, while variance decreases considerably. With respect to age, both variance and bias are much larger at the ends of the age range, even with large sample sizes. The same pattern is replicated for each percentile curve, although the 1^*st*^ percentile is the only one to show noticeable negative bias, associated with not curving high enough in the middle of the arch.

**Figure 6:**
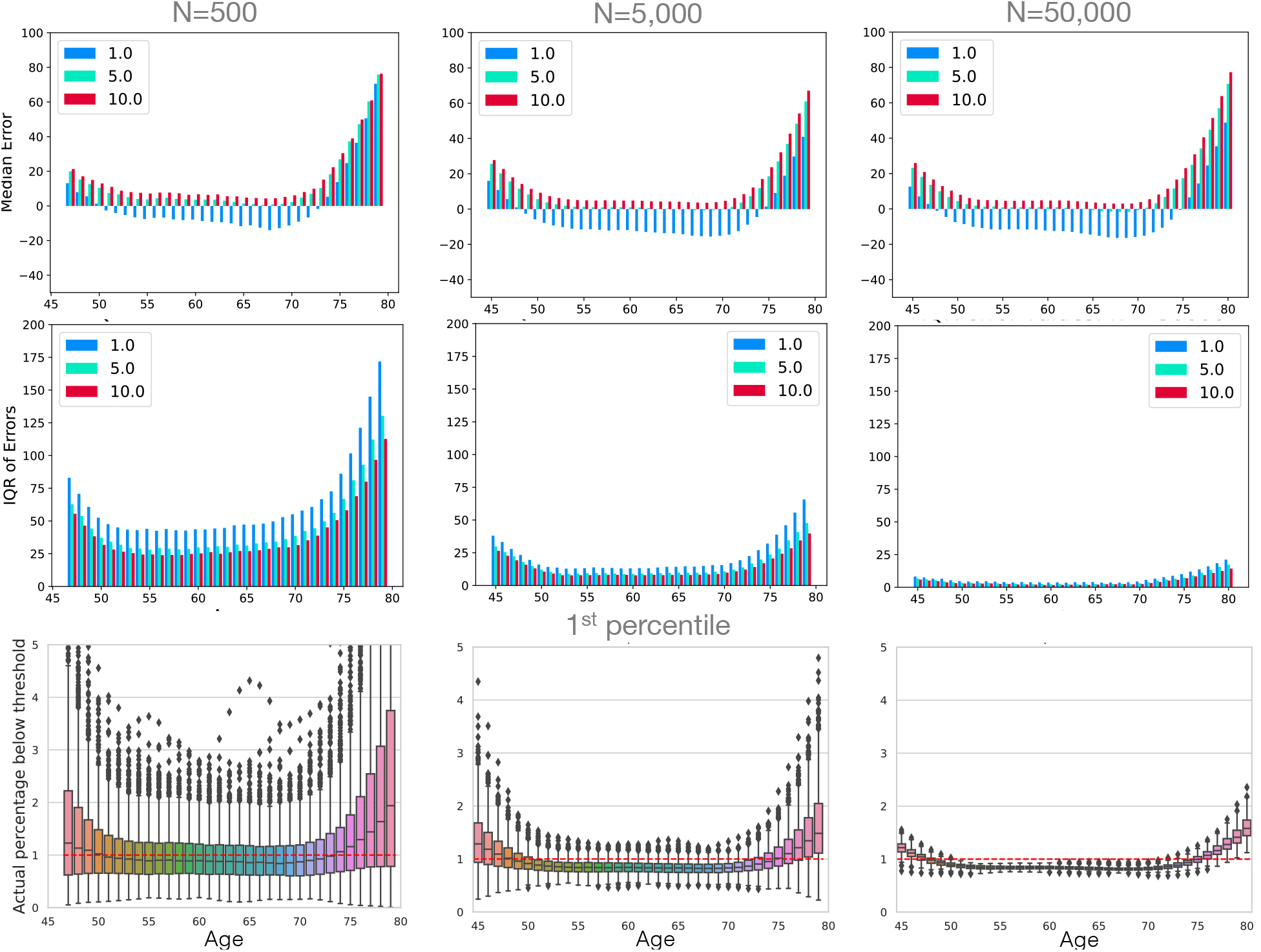
Evaluation of fitting uncertainty with respect to age and sample sizes (columns) 500, 5,000 and 50,000 using GAMLSS with SHASH-*μ*-*σ* and the nonlinear ground truth with non-constant variance. The top row shows the fitting bias, quantified by median error (*E*_1_), for the 1^*st*^ (blue), 5^*th*^ (green) and 10^*th*^ (red) percentiles. The middle row shows the variance of the fitting, measured by interquartile range IQR of errors (*E*_1_). The bottom row shows the actual estimated 1^*st*^ percentiles and the correct percentile value (red dotted line). Results for the 5^*th*^ and 10^*th*^ percentiles are reported in supplementary figure S4 and further sample sizes are reported in supplementary figure S5.

The bottom row in Figure 6 shows the estimated percentile values using boxplots, only for the 1^*st*^ percentile curve in this case. These values represent the expected percentage of normal participants that would have a value under the estimated percentile curve. The error, *E*_2_, is equal to the difference between this value and the nominal percentile (*p* = 1).

These values have a more intuitive or direct connection to applications where the percentile curves are used to identify individuals as having, or being at risk of, some form of pathology. That is, when estimating the 1^*st*^ percentile curve we would expect to get 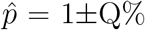, and it is exactly the quantity 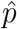 that the boxplots display, where Q (or IQR, a measure of the variation in the results) can be read off the box height (i.e. Q is half the height of the box) and the centre of the boxplots represents the median value, with any bias shown by deviations of this from the correct value (*p* = 1).

For the case where the sample size is 500 data points (participants), the results are very poor, with the variation (value of Q) around 0.5% in the middle of the range and 2-3% at the ends. The whiskers of the boxplots are also important to consider, as half of all results will lie outside of the box (and so is closer to a 95% confidence interval). In this case the range of the whiskers is rarely ever below 2% and becomes extremely large at the ends. For 5,000 data points, which is larger than that used in a number of nomograms in the neuroimaging literature, the value of Q is around 0.2% for much of the age range, but gets close to 1% at the ends of the age range. Furthermore, the range of the whiskers is around 1% in the middle of the age range and reach over 2% at the ends of the range, which could be problematic for some applications. Even with a very large sample size (50,000 data points) the size of the whiskers is near 1% at the ends of the age range, though much smaller in the middle where the variation becomes very small and exposes a bias that is larger than the variation. Boxplots for some of the other sample sizes and other percentiles can be found in the supplementary material in Figures S4 and S5.

## 4. Discussion

Normative modelling has been emerging in neuroimaging in recent years with the availability of big data. From the results shown in this study we want to raise caution when using small or moderate sample sizes for normative datasets. Furthermore, we want to highlight the importance of considering and measuring both variance and bias when evaluating a model, which may not be evident when analysing model performance using a single metric.

We generated samples of different sizes, ranging from 50 to 50,000 data points, consisting of simulated hippocampal volumes for individuals with ages between 45 and 80 years. This age distribution resembled the one in the UK Biobank in order to simulate a real case scenario. We assessed several normative modelling methods, based on sliding window methods or GAMLSS with various settings, using simulated data, with both linear and nonlinear functions.

The results across all fitting methods that we implemented were generally comparable, although GAMLSS with the SHASH transformation showed slightly better overall performance (Figures 2 and 3). This was in line with the recent findings and recommendations by Dinga et al. (2021). Our results show that using these more flexible estimation models is beneficial, particularly with larger samples or cases where the ground truth has a non-negligible nonlinear component. As expected, it is evident that linear models alone are not good for highly nonlinear ground truth distributions, although they demonstrate low variance and hence high repeatability that might make them appear to be working well. Since these linear models do not outperform the more flexible models in the linear ground truth case by very much, we would not recommend the use of purely linear fitting models for normative modelling.

Our results show that a precise estimation of percentiles requires a large sample. Datasets with less than 5000 data points are unlikely to provide accurate estimates for outlying percentiles across the age range (see Figures 2, 3, 5, 6), even though they are not uncommon in the literature. For most sample sizes the variance was dominant, whilst for large sample sizes (e.g. *N*_*s*_ = 50, 000) the bias tended to dominate, making further increases in sample size less useful in improving performance compared to choosing an appropriately flexible method to minimise the bias, although it should be noted that the bias itself was small.

Many studies and datasets have a relatively small number of participants at the ends of the age range, as in the UK Biobank on which we based our simulated age distribution. The substantial deterioration of the performance at the ends of the age range is likely to be due in large part to this and also in part to the difficulty of constraining flexible models at the end of their range. These issues have also been noted in Fraza et al. (2021); Dinga et al. (2021) where they observed bigger deviations in Gaussian-based models in the tails of the distribution, which contained a relatively small proportion of the data. A further contributing factor in real data sets is that the ground truth may be more dynamic at the ends of the human lifespan as well as more varied across individuals. Whatever the cause, the uncertainty at either end of the age range means that extrapolating values beyond the captured age range is likely to be extremely poor and normative models should not be trusted to do this unless the normative sample is extremely large (Figures 4, 5 and 6).

In practice it is usual that only one set of data points is available and the ground truth is unknown, in which case the entire range of possible percentiles shown in the boxplots (in Figures 6, S4 and S5) should be considered, since half of all estimations lie outside the central interquartile range. This would mean that with 5000 participants the estimated 1^*st*^ percentile curve could be closer to the real percentiles in the range 0.2% to 3%, especially near the ends of the age range. If this was used to screen individuals then it would mean that instead of 1% of normal individuals being labelled as positives (those below the curve) this percentage might actually be as high as 3% (or as low as 0.2%) for those near the ends of the age range for an estimated normative model. The consequence for those participants that actually had a pathology would either be beneficial (a better chance of being detected if the estimated percentile curve was high) or, more problematically, detrimental (less chance of detection if the estimated percentile curve was low). To quantify this in terms of statistical power would require knowledge of the distribution of the main quantity (e.g. hippocampal volume) for participants with pathology, though estimated percentile curves that are lower than the true percentile curve will be likely to lead to high false negative rates.

Recent works have modelled different neuroimaging-derived measures using large datasets providing normative or reference curves across the lifespan (Bethlehem et al., 2022; Rutherford et al., 2022). Our work complements these by focusing on evaluating performance of commonly used normative modelling methods using simulated samples. Using these we have evaluated the performance across multiple options, showing that summarising over ages can hide poor results and representing the performance as a single metric is likely to be too simplistic. Furthermore, measuring only variance (e.g. using IQR) is not sufficient to judge performance and define an appropriate fitting method. Thus, it is recommended to either assess bias independently from variance, as we have seen very different behaviour of these two terms as the sample size is increased. However, exact bias measurements can only be made if the ground truth is known, and empirically estimating it reliably requires extremely large samples (as outlined in the methods section) especially since near the end of the age range the number of data points is likely to be smaller. Therefore, using a simulation-based approach can be a very useful method for assessing performance, and for this reason we have made our code available as a general resource for evaluation of normative modelling methods.

There are several limitations of our work that should be considered. One limitation is the sole use of simulated data. We decided to focus on simulations for several reasons. Firstly, because we wanted to apply and evaluate models on large sample sizes (up to 50,000 data points). With real data this would only be possible by merging datasets from different sites and/or studies (e.g. Thompson et al. (2014); Casey et al. (2018); Bethlehem et al. (2022)). This would then require data harmonisation, which represents a separate issue and active field of research on its own, as the application of normative models and normative ranges in real datasets should be adjusted to effectively deal with site-effects (Kia et al., 2020, 2021). To make our simulations realistic we used an age distribution based on the UK Biobank, the biggest single-study dataset currently available, which has been already used in several normative modelling studies (Nobis et al., 2019; Janahi et al., 2021). This makes our simulations easier to compare with these studies, with the added advantage of knowing the ground truth distribution to give a sense of how good the estimations are in the real dataset studies. Another limitation is the choice of ground truth distributions, using only two cases: the simplest one of a completely linear function with constant variance and perfectly Gaussian conditional distributions. We did experiment with other distributions, but we believe that these two showed all the interesting effects and spanned a reasonable range of expected true distributions, although still being slightly idealised by the use of Gaussian conditionals. However, by making the ground truth somewhat simpler than might occur in practice we are creating a situation where the normative models are likely to do slightly better than they might do otherwise. Consequently, these results are likely to be slightly better than could be expected in reality, acting like an upper bound on performance. This argues even more strongly for applying normative modelling only with very large samples.

This approach (code available at https://github.com/jelenabozek/NormativeModelling) can be easily extended to work with different data distributions, and used to evaluate the performance of a normative model even before collecting data. Given a hypothesis on the distribution of the values and the expected/known distribution of data points in the age bins, the simulations can provide an estimate of the accuracy of the percentiles of interest in terms of both variance and bias. This can, in turn, inform power calculations (e.g., how many participants are needed to reach a certain level of accuracy in estimating the 5^*th*^ percentile for people over 70 years old?) and decisions required for applications in a clinical context (e.g., given a certain normative model, how reliable would a cutoff based on the 5^*th*^ percentile be for people over 70 years old?).

## 5. Conclusion

Normative modelling of simulated values (e.g. hippocampal volumes) from samples with sizes ranging from 50 to 50,000 confirmed that flexible models perform better (e.g. GAMLSS with SHASH *μ*-*σ* transformation), especially when the ground truth is nonlinear. Surprisingly large samples with several thousand data points are needed to accurately capture outlying percentiles across the age range for applications in research and clinical settings. Assessment of the reliability of the model’s estimation of the percentiles is important for the clinical setting and should be carefully considered. Summarising evaluation results into a single summary value would often not be sufficient for assessing performance, especially if it did not include some part that was sensitive to bias, when selecting appropriate fitting methods. Thus, both bias and variance, or something sensitive to both, should be used when assessing the model’s performance. Furthermore, extreme caution is needed when attempting to extrapolate beyond the age range included in the source dataset, due to the rapid increase in uncertainty at the ends of the age range. To help with such evaluations of normative models we have made our code available and encourage researchers to use this when developing or utilising normative models.

## 6. Data/code availability statement

The code used for generating the simulated data, fitting the normative models and evaluating the models performance is openly available at https://github.com/jelenabozek/NormativeModelling.

## 7. Acknowledgements

LG is supported by an Alzheimer’s Association Grant (AARF-21-846366) and by the National Institute for Health and Care Research (NIHR) Oxford Health Biomedical Research Centre (BRC). MJ is supported by the NIHR Oxford Biomedical Research Centre (BRC), and this research was funded by the Wellcome Trust (215573/Z/19/Z). This work was also supported by the Wellcome Centre for Integrative Neuroimaging, which has core funding from the Wellcome Trust (203139/Z/16/Z). For the purpose of open access, the authors have applied a CC-BY public copyright licence to any Author Accepted Manuscript version arising from this submission.

## 8. Declaration of Competing Interest

The authors declare that they have no known competing financial interests or personal relationships that could have appeared to influence the work reported in this paper.

## Appendix A.

Supplementary Material

**Figure S1:**
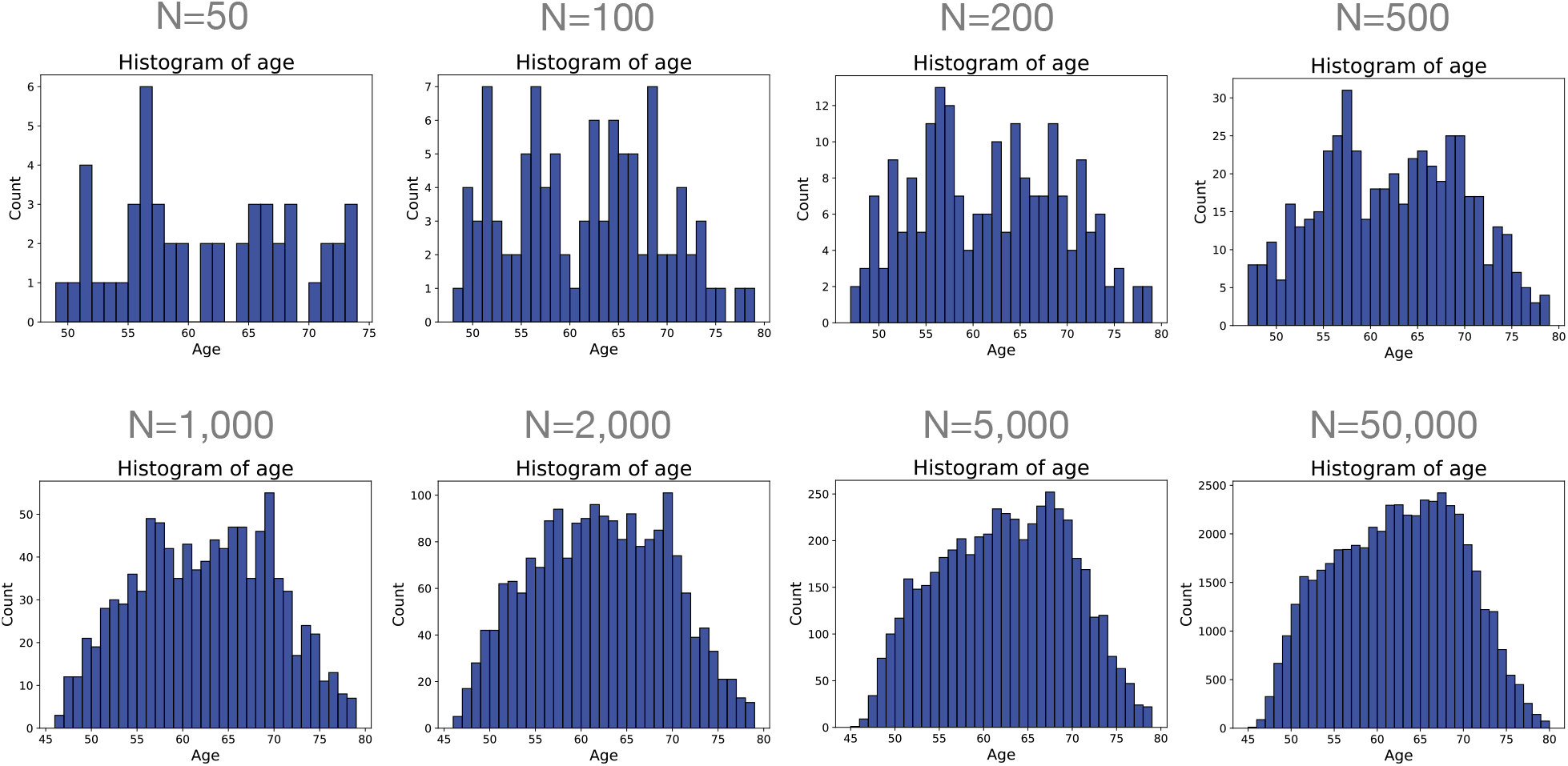
Histograms of ages for the samples used in the simulations. The number of datapoints is shown in the figure and all simulations with the same number of datapoints use the same set of ages, hence why there is one fixed histogram for each N value.

**Figure S2:**
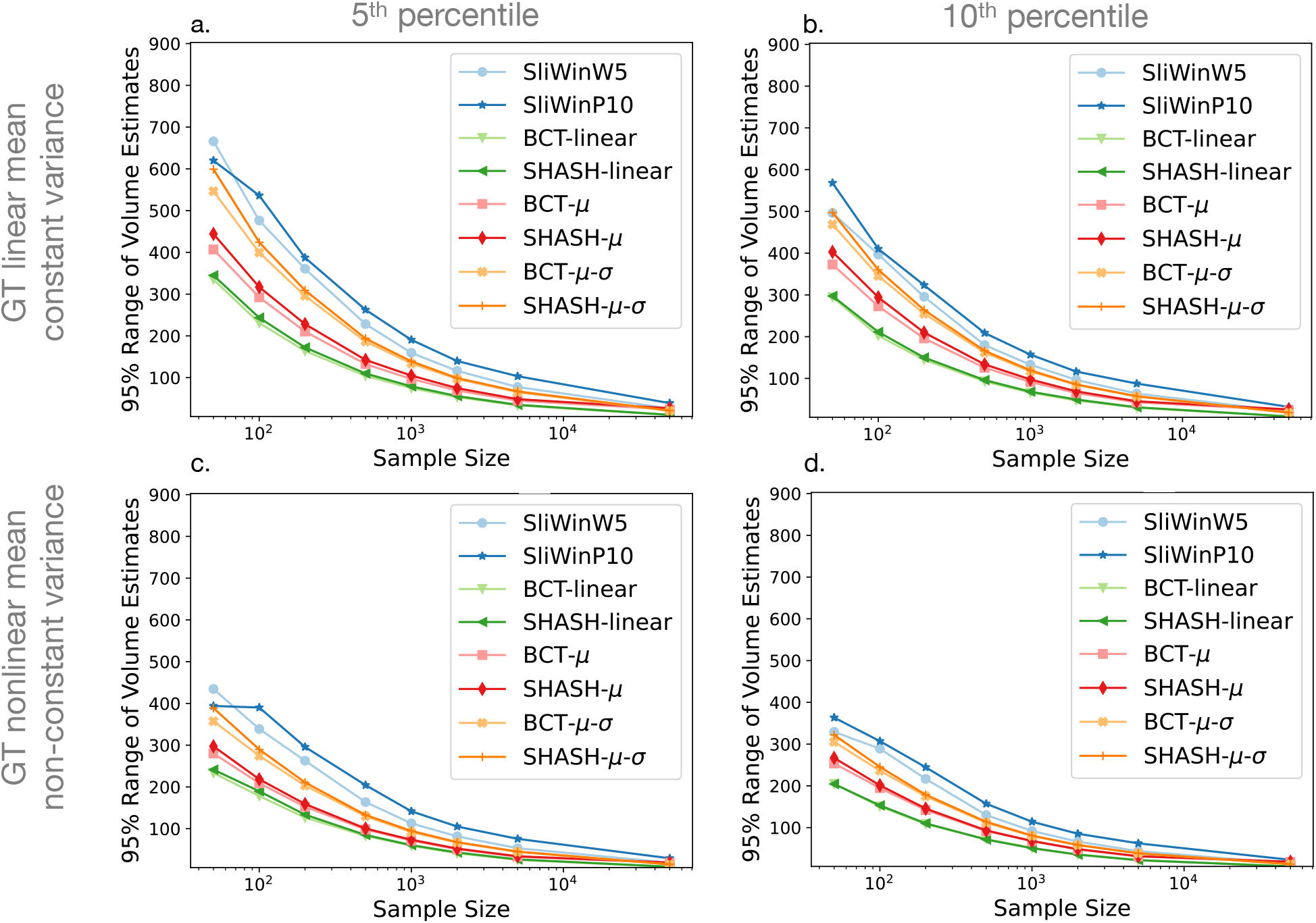
95% range of volume estimates against sample size for each fitting method. The plots show the confidence intervals (median across age) of the 5^*th*^ percentile (left column) and 10^*th*^ percentile (right column) curves for linear mean and constant variance ground truth (top row) and nonlinear mean and non-constant variance ground truth (bottom row). Results for the 10^*th*^ percentile are very similar to those for the 5^*th*^ percentile (also reported in Figure 2 of the main text).

**Figure S3:**
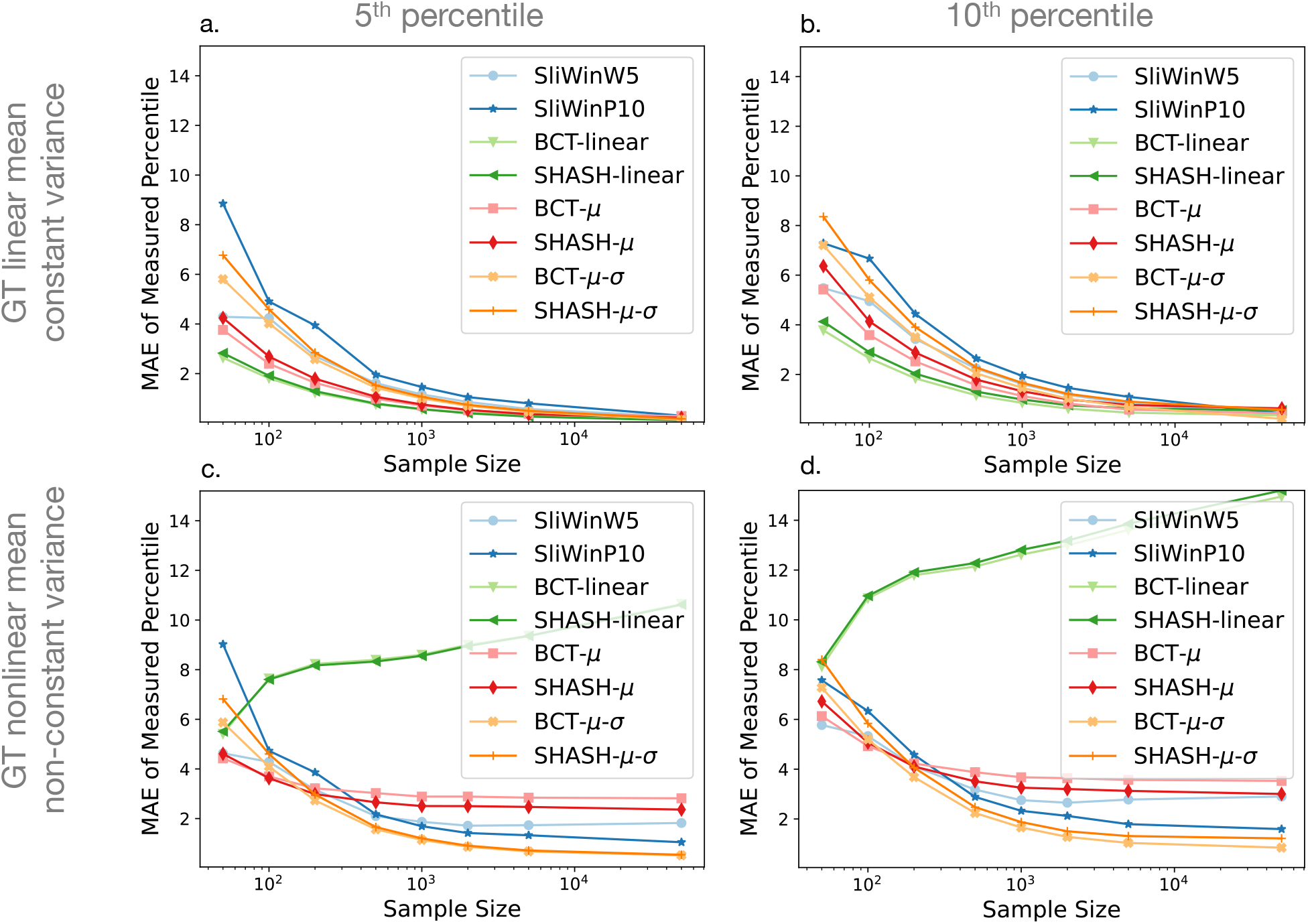
Mean Absolute Error (MAE) against sample size for each fitting method. The plots show the mean across age of the mean absolute error of the estimates of percentile error (*E*_2_) of the 5^*th*^ percentile (left column) and 10^*th*^ percentile (right column) curves for linear mean and constant variance ground truth (top row) and nonlinear mean and non-constant variance ground truth (bottom row). Results for the 10^*th*^ percentile are very similar to those for the 5^*th*^ percentile (also reported in Figure 3 of the main text).

**Figure S4:**
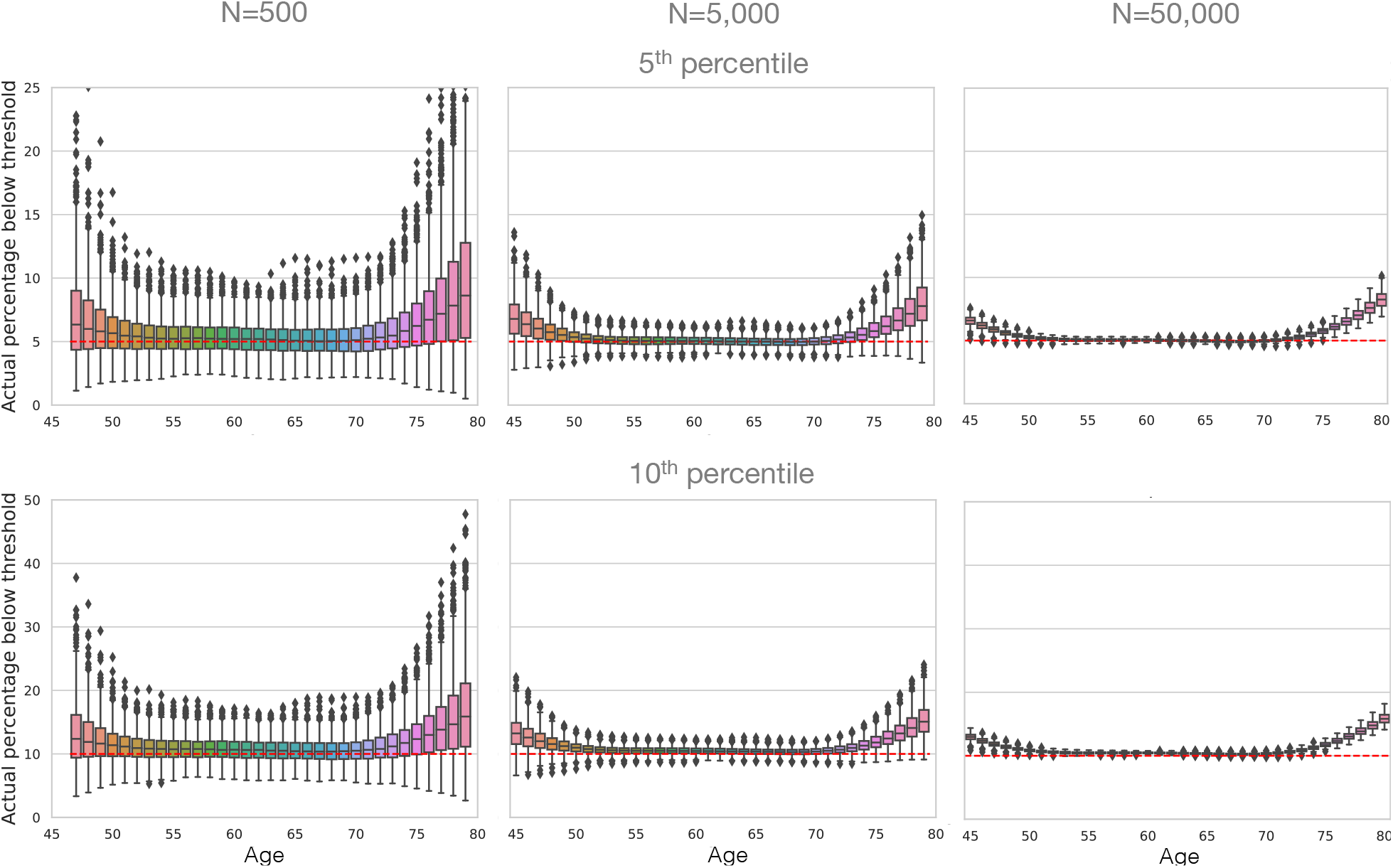
Evaluation of fitting uncertainty with respect to age for sample sizes (columns) 500, 5,000 and 50,000 using GAMLSS with SHASH-*μ*-*σ* and the nonlinear ground truth with non-constant variance. Actual estimated 5^*th*^ (top row) or 10^*th*^ (bottom row) percentiles and the correct percentile value (red dotted line) are shown. Results for 1^*st*^ percentile are reported in Figure 6 of the main text.

**Figure S5:**
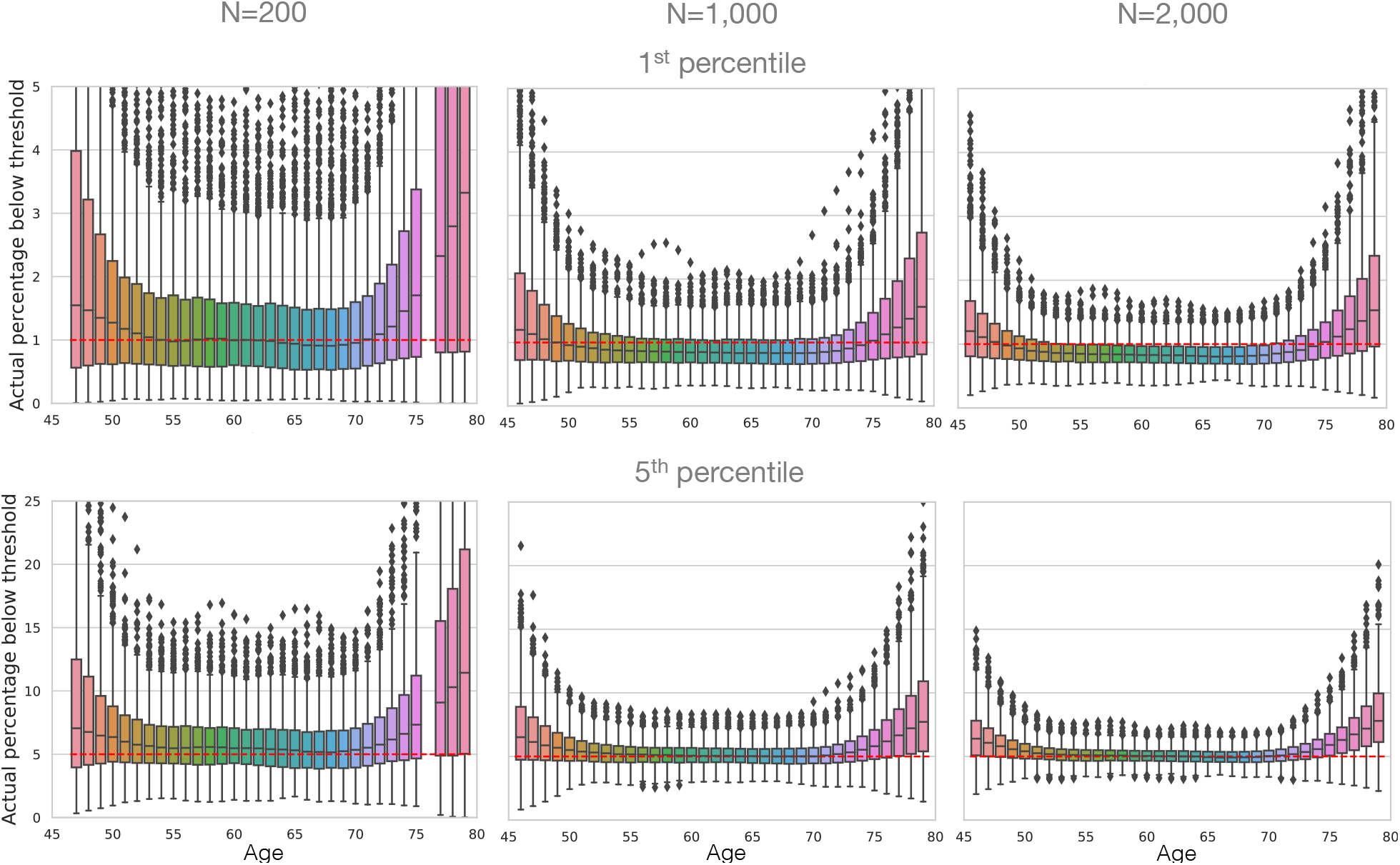
Evaluation of fitting uncertainty with respect to age for sample sizes (columns) 200, 1,000 and 2,000 using GAMLSS with SHASH-*μ*-*σ* and the nonlinear ground truth with non-constant variance. Actual estimated 1^*st*^ (top row) or 5^*th*^ (bottom row) percentiles and the correct percentile value (red dotted line) are shown. Results for other sample sizes are reported in Figure 6 and Figure S4.

## Notes

### Competing Interest Statement

The authors have declared no competing interest.

